# VST Family Proteins are Regulators of Root System Architecture in Rice and *Arabidopsis*

**DOI:** 10.1101/2020.05.13.091942

**Authors:** Yanlin Shao, Kevin R. Lehner, Hongzhu Zhou, Isaiah Taylor, Chuanzao Mao, Philip N. Benfey

## Abstract

Root System Architecture (RSA) is a key factor in the efficiency of nutrient capture and water uptake in plants. Understanding the genetic control of RSA will be useful in minimizing fertilizer and water usage in agricultural cropping systems. Using a hydroponic screen and a gel-based imaging system we identified a rice gene, *OsVST1*, which plays a key role in controlling RSA. This gene encodes a homolog of the *Arabidopsis* VAP-RELATED SUPPRESSORS OF TMM (VSTs), a class of proteins that promote signaling in stomata by mediating plasma membrane-endoplasmic reticulum contacts. *OsVST1* mutants have shorter primary roots, decreased root meristem size, and a more compact root system architecture. We show that the *Arabidopsis* VST triple mutants have similar phenotypes, with reduced primary root growth and smaller root meristems. Expression of *OsVST1* largely complements the short root length and reduced plant height in the *Arabidopsis* triple mutant, supporting conservation of function between rice and *Arabidopsis* VST proteins. In a field trial, mutations in *OsVST1* do not adversely affect grain yield, suggesting that modulation of this gene could be used as a way to optimize RSA without an inherent yield penalty.

## Introduction

As living organs, plant roots play crucial roles in water and nutrient acquisition as well as in providing support and anchorage in soil. Root system architecture (RSA), the spatial arrangement of roots for a given plant, reflects these critical functions (Lynch, 1995). In the model dicot, *Arabidopsis thaliana*, RSA is primarily determined by growth rates and the branching patterns of lateral roots (Smith & De Smet, 2012). Asian rice (*Oryza sativa*) and other agronomically important monocots, possess more complex RSA profiles, dominated by the contribution from shoot-derived roots (Hochholdinger & Zimmermann, 2009).

There is strong interest in selecting for specific RSA phenotypes in crops to increase yield and promote stability under stress. Yet, due mainly to the inherent difficulties in measuring largely underground root phenotypes, the genetics underlying RSA have remained elusive. While many different root imaging modalities are being developed, one with an ideal combination of throughput, accuracy, and realistic growth conditions has yet to be produced (Atkinson et al., 2019). In this study, we combined the high-throughput of a hydroponic screening system with the accuracy of a non-invasive, gel-based imaging platform to identify a novel regulator of RSA in rice.

Optimizing RSA for agriculture will require careful consideration of target environments and crop-specific management practices (Bishopp & Lynch, 2015). Depending on soil conditions, fertilizer inputs, and water use, an ideal arrangement of roots may lie anywhere on a continuum from deep and expansive to shallow and dense (Morris et al., 2017). With this in mind, it will be useful to identify the genetics underlying a range of RSA phenotypes.

Despite the challenges of identifying RSA genes in monocots, there have been a few successes. Particularly notable in rice is the cloning of two genes, *DRO1* and *PSTOL1*, that underlie quantitative trait loci for root phenotypes (Gamuyao et al., 2012; Uga et al., 2013). The identification of favorable alleles of these genes has been followed by successful deployment into field trials for increased stress resilience. This suggests the potential utility of harnessing beneficial root architecture traits for yield improvement and stability.

In addition, a number of genes affecting RSA in rice have been identified in genetic screens using mutagenized populations (Mai et al., 2014). Most of these have strong effects on root traits due to the lack of certain root types or other detrimental defects. Many of these mutants also have highly pleiotropic phenotypes. While contributing to our understanding of the genetic requirements of root development, highly reduced grain yield serves as a barrier to the use of these alleles in breeding (Wissuwa et al., 2016).

A common theme of growth regulation in plants is signaling through extracellular receptors, such as receptor-like kinases (RLKs) (De Smet et al., 2009). These are often plasma membrane proteins that bind extracellular ligands and affect downstream transcriptional responses. RLKs in particular are predicted to comprise ~2% of protein coding genes in rice and *Arabidopsis* (Shiu et al., 2004). With such a large diversity of RLKs and downstream proteins, a developing theme is the importance of spatial proximity of protein complexes to properly initiate and propagate a signal transduction cascade (Qi & Torii, 2018).

Recent work uncovered the key roles of a family of *Arabidopsis* genes encoding proteins containing Major Sperm Protein (MSP) domains in stomatal patterning and above-ground plant architecture(Ho et al., 2016). These proteins were found to regulate RLK signaling through establishing or maintaining plasma membrane-endoplasmic reticulum (PM-ER) contact sites and were named VAP-RELATED SUPPRESSORS OF TMM (VSTs). Here we identify genes encoding rice and *Arabidopsis* VSTs as key regulators of RSA.

## Results

### An RSA Mutant with Reduced Root Length and Altered Growth Dynamics

We used a high-throughput hydroponic screening system to identify lines with altered RSA phenotypes within an EMS-mutagenized rice population. We discovered a mutant (designated E104-SR) that had reduced root length compared to the unmutagenized, wild type parent (Fig. S1A and S1B). Additionally, E104-SR had increased crown root number and slight reductions in shoot height at later development stages (Fig. S1C and S1D).

Using a non-destructive, gel-based root imaging system (Galkovskyi et al., 2012; Iyer-Pascuzzi et al., 2010; Topp et al., 2013), we examined differences in RSA between E104-SR and wild-type. At ten days after germination, the mutant had a different RSA profile, characterized by a shorter and more compact root system (Fig. 1A). Imaging at 24-hour intervals across ten days of growth allowed us to monitor the rates of change of RSA traits throughout early development. Network Convex Area is defined as the area of the convex hull encompassing the root system and is a measure of RSA extent (Iyer-Pascuzzi et al., 2010). Major Ellipse Axis is the length of the major axis of the best fitting ellipse to the root system and is a proxy for RSA depth (Galkovskyi et al., 2012). Both Network Convex Area and Major Ellipse Axis increase throughout the ten-day time-course in wild-type plants, while the E104-SR mutant shows a gradual flattening in the rates of change of these traits (Fig. 1B and C).

**Figure 1:**
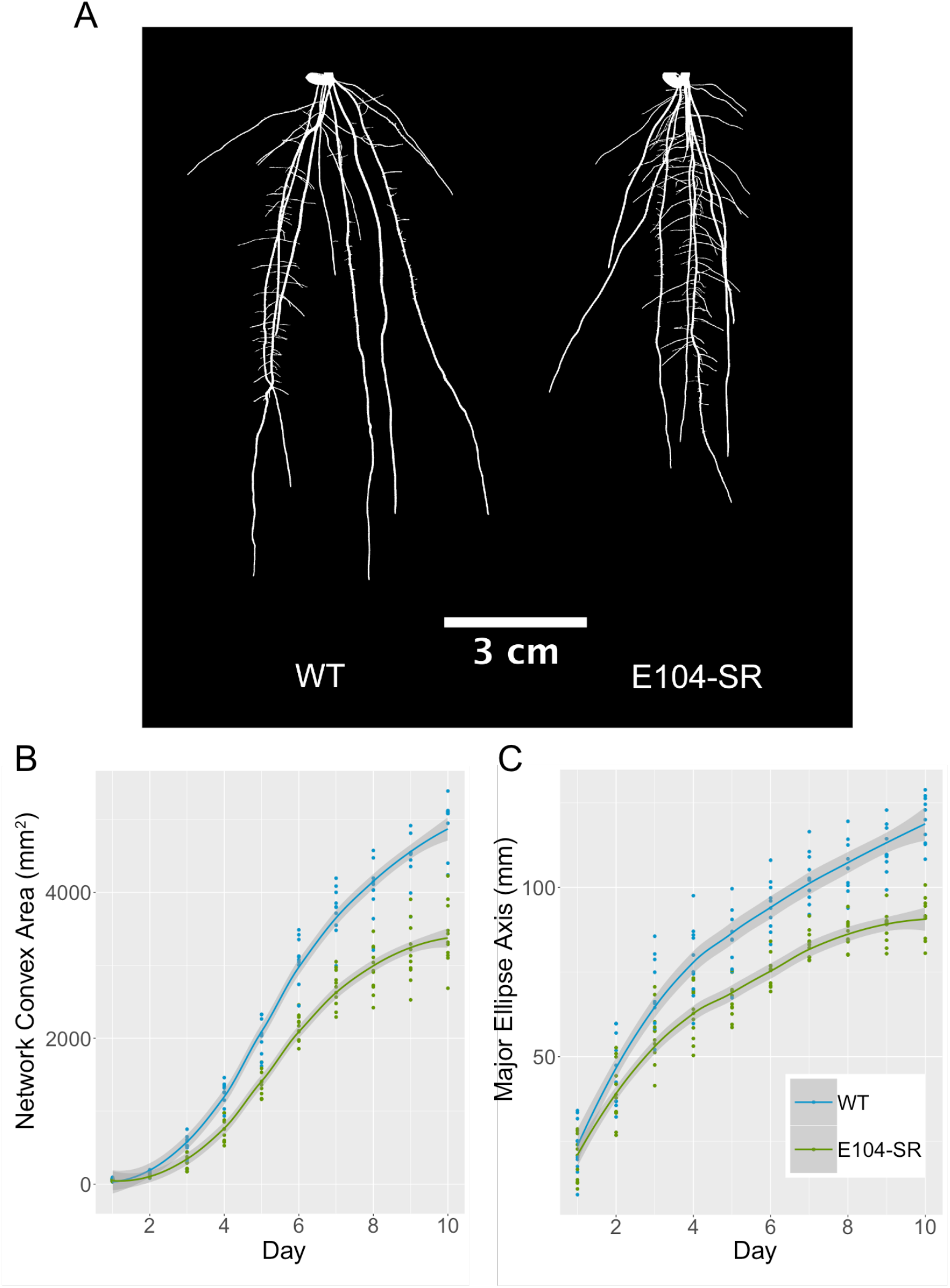
E104-SR mutant plants have shorter, more compact root systems and altered growth dynamics. (A) Representative thresholded images of 10-day-old, gel-grown, roots. (B) Network Convex Area of wild-type and E104-SR roots throughout ten days of imaging. (C) Major Ellipse Axis of wild-type and E104-SR roots throughout ten days of imaging. Line is fitted with the “geom_smooth” function in ggplot2.

### E104-SR Roots have Smaller Meristems

To better understand the nature of the E104-SR mutant phenotype, we examined longitudinal sections of E104-SR and wild-type primary roots. The mutant roots have significantly smaller meristematic regions during early growth (Fig 2A-2C). Additionally, primary root width is reduced in mutants (Fig 2D).

**Figure 2:**
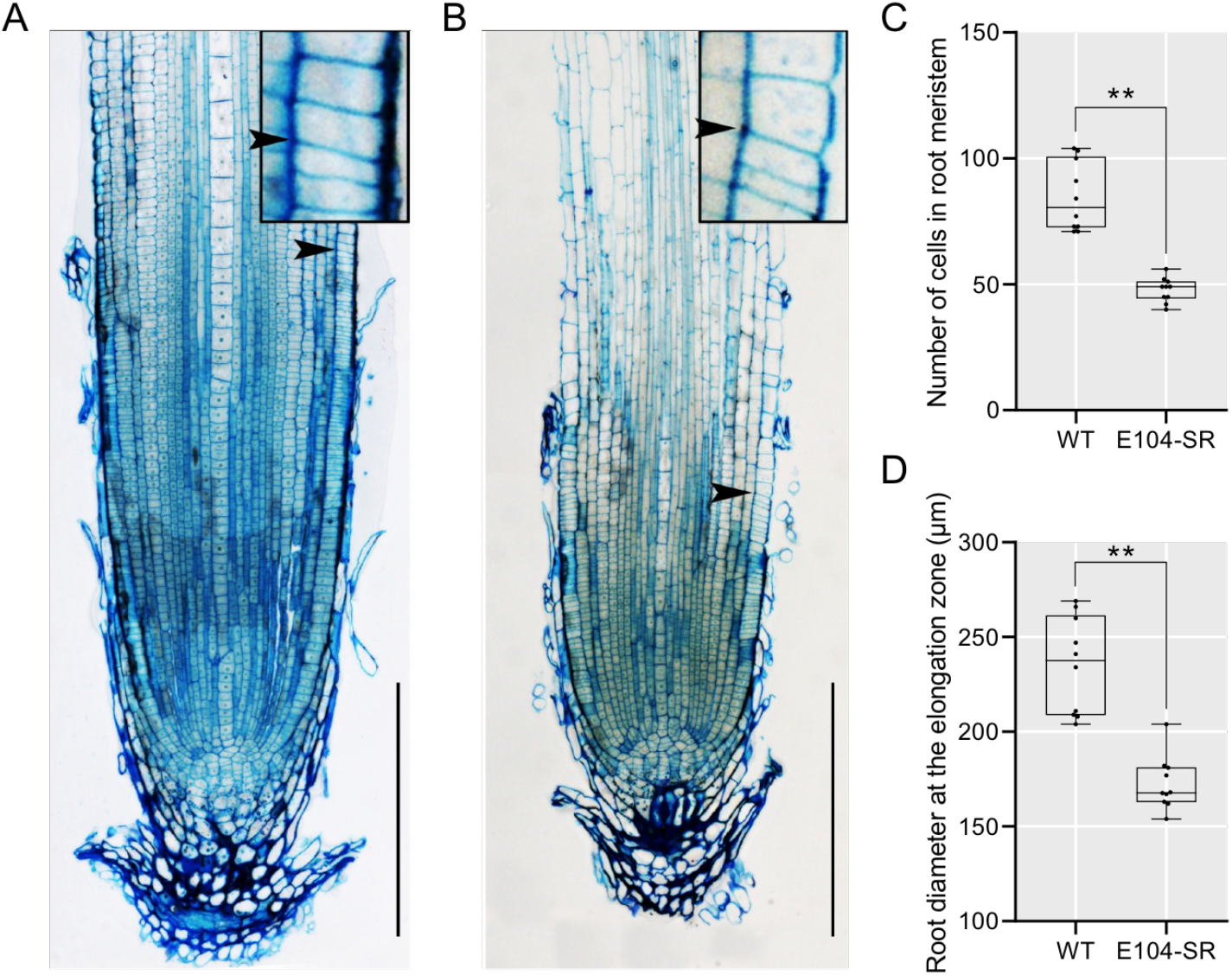
E104-SR has reduced root meristem size and decreased root width. (A, B) Longitudinal sections of the primary root of 5-day-old seedlings of wild-type (A) and E104-SR (B). Arrow indicates location of junction of meristem and elongation zone. Bars = 200 μm. (C, D) Quantification of root meristem cell number (C) and root diameter at the elongation zone (D). Median is represented by horizontal line in the box plot, box ranges represent quartiles 1 and 3, and minimum and maximum values are represented by error bars (*n* = 10). Data significantly different from the corresponding wild type are indicated (**P < 0.01; Student’s t test).

### The Causative Mutation in E104-SR Maps to an 11-kilobase Deletion

To identify the mutation underlying the mutant RSA phenotype, we used a mapping-by-sequencing approach based on bulked segregant analysis (Michelmore et al., 1991). We backcrossed the E104-SR mutant to the unmutagenized parental cultivar and selfed the F_1_ to generate a BC_1_F_2_ mapping population. Using our gel imaging platform, we phenotyped ~120 BC_1_F_2_ plants, isolating DNA from 20 plants with the deepest and 20 plants with the most shallow root systems. Each of these groups of DNA samples was pooled separately, making up the mutant and wild-type bulks, respectively.

We performed paired-end, short read sequencing on the two contrasting bulks and used the R package QTLseqr (Mansfeld & Grumet, 2018), with an approach similar in concept to MUT-map (Abe et al., 2012). This analysis identified a candidate region on chromosome 8 that contained the likely causative mutation underlying the E104-SR root phenotype (Fig S2). Further genotyping of the individuals that comprised the mutant bulk allowed for fine-mapping of the causative region to an interval between approximately 2.2-3.5 Mb on chromosome 8.

### Deletion of a Single Gene is Responsible for the E104-SR Phenotype

Although no EMS-induced SNPs were found within the interval on Chromosome 8 between 2.2-3.5 Mb, there was an 11-kilobase deletion located at 2.3 Mb that was highly enriched in the mutant bulk (Fig 3A). The deletion results in the complete loss of two genes. The first, *LOC_Os08g04620*, is predicted to encode a Major Sperm Protein (MSP) domain containing protein. The second, *LOC_Os08g04630*, is annotated as encoding a mitochondrial-localized protein, an external NADH-ubiquinone oxidoreductase. Both of these genes are expressed in roots, and at higher levels than in shoots (Raines et al., 2016).

**Figure 3:**
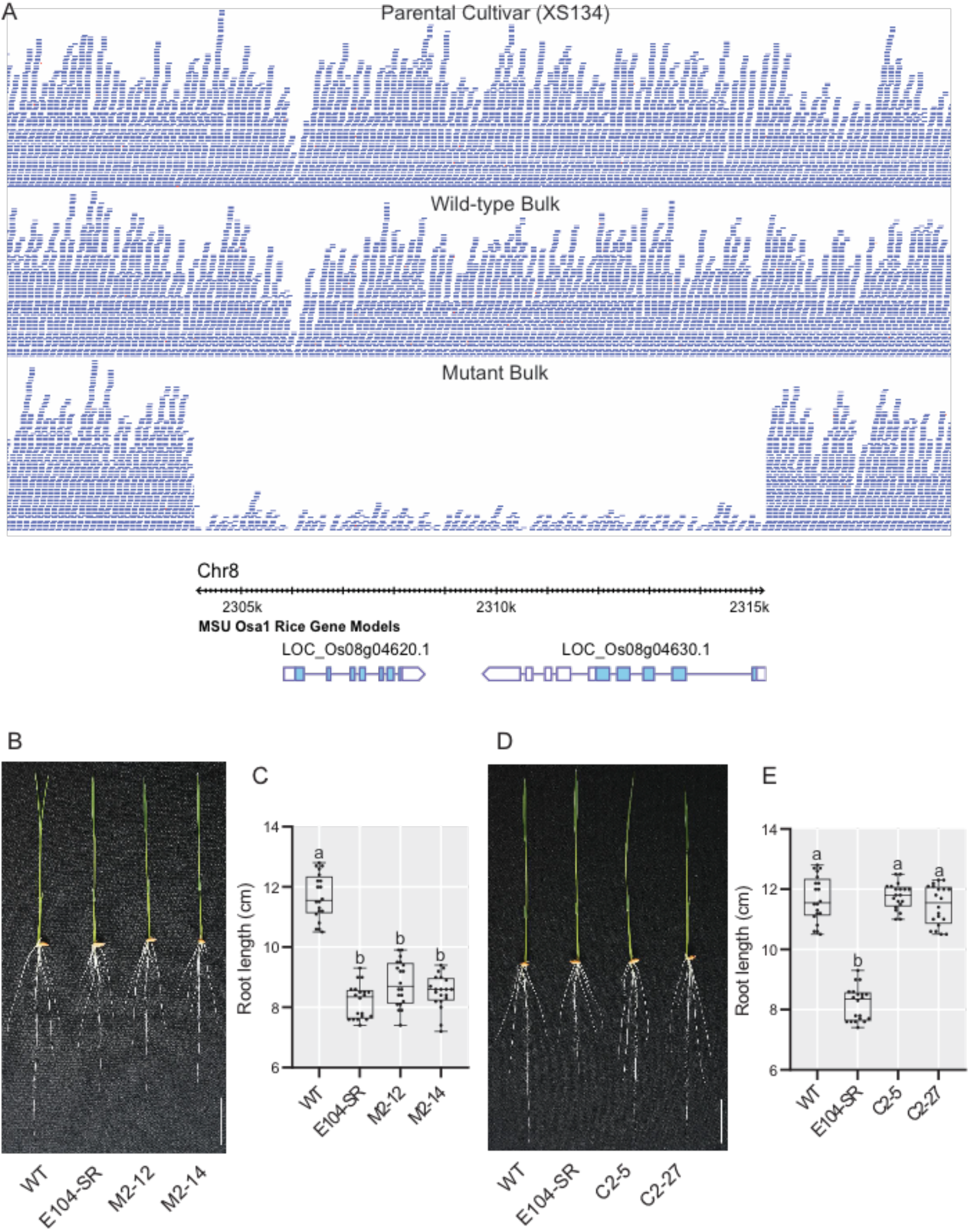
*LOC_Os08g04620* is located within a deletion in E104-SR and is the causative gene for the reduced root length phenotype. (A) Sequence read depth from the parental cultivar, wild type, and mutant bulks with the MSU v7 gene models within the deleted region shown below. (B, C) Phenotypes (B) and root lengths (C) of 7-day-old knockout transgenic lines grown in hydroponic culture. M2-12 and M2-14 are two independent CRISPR-Cas9 generated *LOC_Os08g04620* knockout T1 lines. (D, E) Phenotypes (D) and root lengths (E) of 7-day-old complementation lines grown in hydroponic culture. C2-5 and C2-27 are two independent T2 transgenic lines of E104-SR containing a genomic fragment with only *LOC_Os08g04620*. Bars = 3 cm. Median is represented by horizontal line in the box plot, quartiles 1 and 3 are represented by box ranges, and minimum and maximum values are represented by error bars (*n* = 20). Significantly different values are indicated by different letters (P < 0.01; one-way ANOVA with Tukey’s test).

To determine which of these candidates is responsible for the E104-SR root phenotype, we took two complementary approaches. First, we generated additional loss-of-function alleles of *LOC_Os08g04620* and *LOC_Os08g04630* using CRISPR-Cas9 (Fig. S3 and S4). In hydroponics, two independent mutant alleles of *LOC_Os08g04620* (M2-12 and M2-14) had root lengths that are shorter than the wild-type parent and indistinguishable from that of E104-SR (Fig 3B and 3C). In contrast, plants with two independent mutant alleles of *LOC_Os08g04630* (M3-6 and M3-7) had root lengths similar to wild type (Fig S5). Next, we constructed transgenic lines with genomic fragments containing either *LOC_Os08g04620* or *LOC_Os08g04630* in the E104-SR mutant background to test for complementation of the reduced root length phenotype. Two lines with *LOC_Os08g04620* (C2-5 and C2-27) had root lengths comparable to wild type (Fig 3D and 3E). Lines containing *LOC_Os08g04630* (C3-3 and C3-5), however, failed to complement the mutant phenotype (Fig. S6). These data support the conclusion that deletion of *LOC_Os08g04620* is responsible for the reduced root length phenotype in E104-SR.

### *LOC_Os08g04620* Encodes a Homolog of the *Arabidopsis* VST Proteins

*LOC_Os08g04620* encodes a protein that has a domain architecture characterized by a central MSP domain and a C-terminal coiled-coil domain (Fig S7B). The protein has a predicted nuclear localization sequence, but lacks a transmembrane domain. This structure is similar to a recently characterized family of proteins in *Arabidopsis*, the VAP-RELATED SUPPRESSORS OF TMM (VSTs) (Ho et al., 2016). The *Arabidopsis* VST proteins were identified as key regulators of stomatal patterning. They were shown to function as peripheral plasma membrane proteins, promoting PM-ER contacts to facilitate signaling. Consistent with this, despite not having any predicted transmembrane domains, the rice protein encoded by *LOC_Os08g04620* was detected as PM-localized in a proteomics study (Natera et al., 2008). In a multiple sequence alignment of MSP domain-containing proteins in *Arabidopsis* and rice, LOC_Os08g04620 clusters with the three *Arabidopsis* VST proteins and an apparent rice paralog (LOC_Os07g37270) (Fig S7A). The rice genes *LOC_Os08g04620* and *LOC_Os07g37270* have distinct organ-level expression profiles, with *LOC_Os08g04620* notably being more highly expressed in roots than *LOC_Os07g37270* (Raines et al., 2016). As *LOC_Os08g04620* and *LOC_Os07g37270* encode proteins that have high degrees of sequence similarity with the *Arabidopsis* VSTs, we will refer to them as *OsVST1* and *OsVST2*, respectively.

### *OsVST1* is expressed broadly in roots

We assessed expression of *OsVST1* using a line containing a transcriptional reporter with the *OsVST1* promoter fused to the beta-glucuronidase gene (*GUS*). *OsVST1* is expressed broadly throughout the root, including primary root, crown roots, and lateral roots (Fig 4A and 4B). *OsVST1* was expressed in the epidermis, exodermis, sclerenchyma, and stele of the root maturation zone (Fig 4C). *OsVST1* also showed high levels of staining at the stem base (the region of crown root initiation) (Fig 4D).

**Figure 4:**
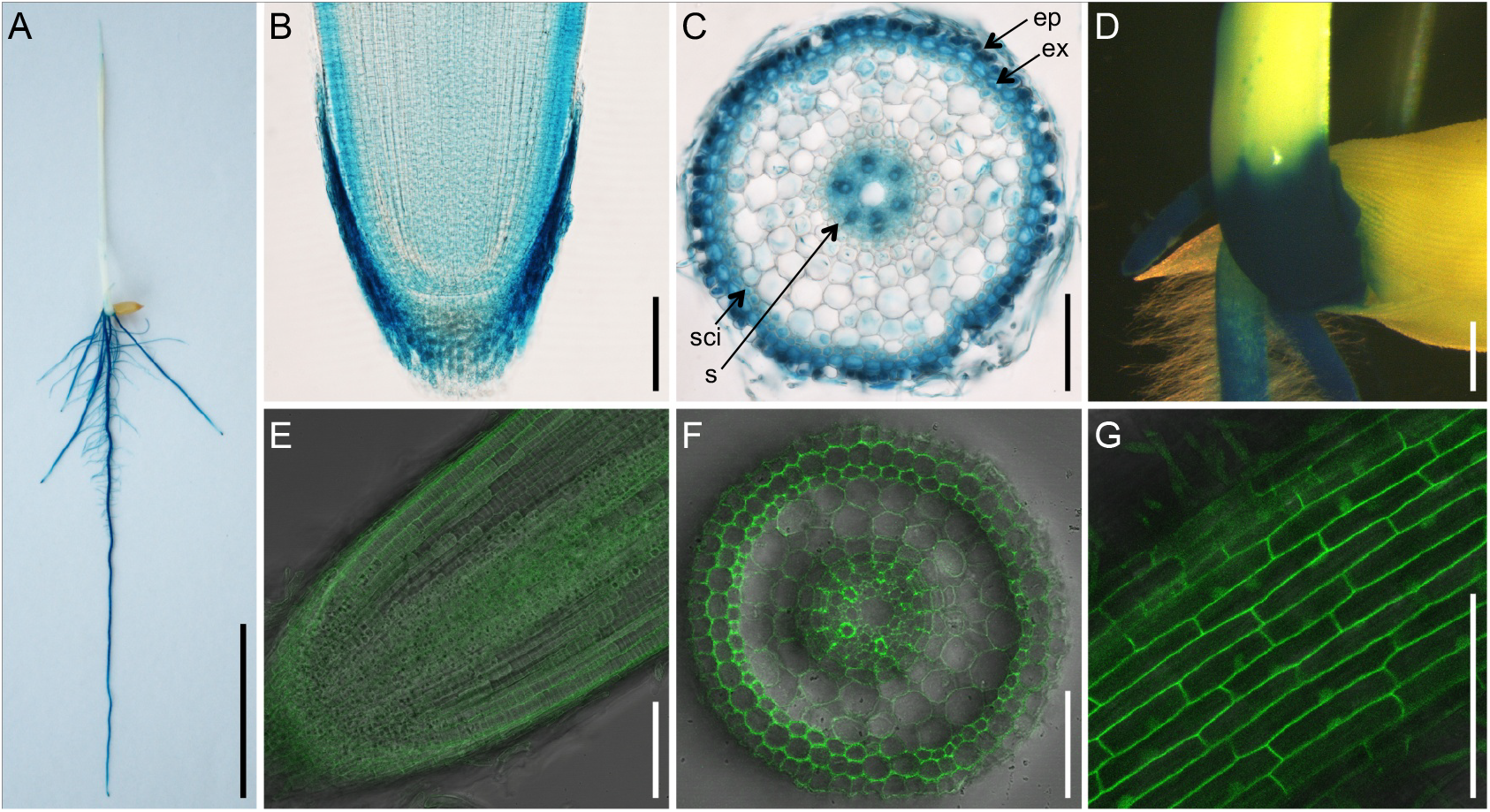
Expression and protein accumulation pattern of OsVST1 are consistent with mutant RSA phenotype of E104-SR. (A) to (D) GUS staining of five-day-old *ProVST1:GUS* transgenic seedlings. Whole seedling (A), longitudinal section of the root tip (B), cross section of the primary root maturation zone (C), and stem base (D). (E) and (F) Tissue specificity of accumulation pattern of GFP-VST1 protein in longitudinal section of root tip (E) and cross section of primary root maturation zone (F). (G) Subcellular localization of OsVST1 protein in cortex cell of root. Abbreviations: ep, epidermis; ex, exodermis; scl, sclerenchyma; s, stele. Bars, 3 cm in (A); 100 μm in (B), (C), (E), (F), and (G); 1 mm in (D).

To analyze patterns of OsVST1 protein accumulation and subcellular localization, we produced transgenic plants expressing *OsVST1* fused to either a C-terminal or N-terminal *GFP* (*VST1-GFP* or *GFP-VST1* respectively). Each of these constructs is driven by the *OsVST1* promoter and was transformed into E104-SR mutant plants. Although each of the two transgenic constructs can only partially complement the short root phenotype of E104-SR mutant (Fig S8), the *GFP-VST1* transgenic plants had longer primary roots than the *VST1-GFP* transgenic plants. Therefore, we selected a *GFP-VST1* transgenic plant for further analysis. Using this line, we examined protein localization within the root. We found that OsVST1 protein is present throughout the region near the root tip (Fig 4E). In a cross section of the maturation zone, we detected a high level of GFP fluorescence in the vasculature of the root (Fig 4F). Although OsVST1 mainly appeared to localize to the plasma membrane, we also detected a pattern consistent with nuclear localization in some cells (Fig 4G).

### The *Arabidopsis vst1/2/3* Triple Mutant has Reduced Root Length and Decreased Meristem Size

In the previous characterization of the *Arabidopsis* VST genes, the authors described a compact aboveground growth pattern in the *vst1/2/3* triple mutant (Ho et al., 2016). They showed that expression of *AtVST1* was able to rescue the plant height phenotype. As their analysis did not include any root phenotyping of the *vst1/2/3* mutant, we characterized it and found that primary root length is reduced (Fig.5A). Root length was restored to wild-type levels when *AtVST1* was expressed. Similar to that of the rice line E104-SR, the *vst1/2/3* triple mutant has a shorter root meristem (Fig 5B-C). The reduced meristem size was also rescued by expression of *AtVST1*.

**Figure 5:**
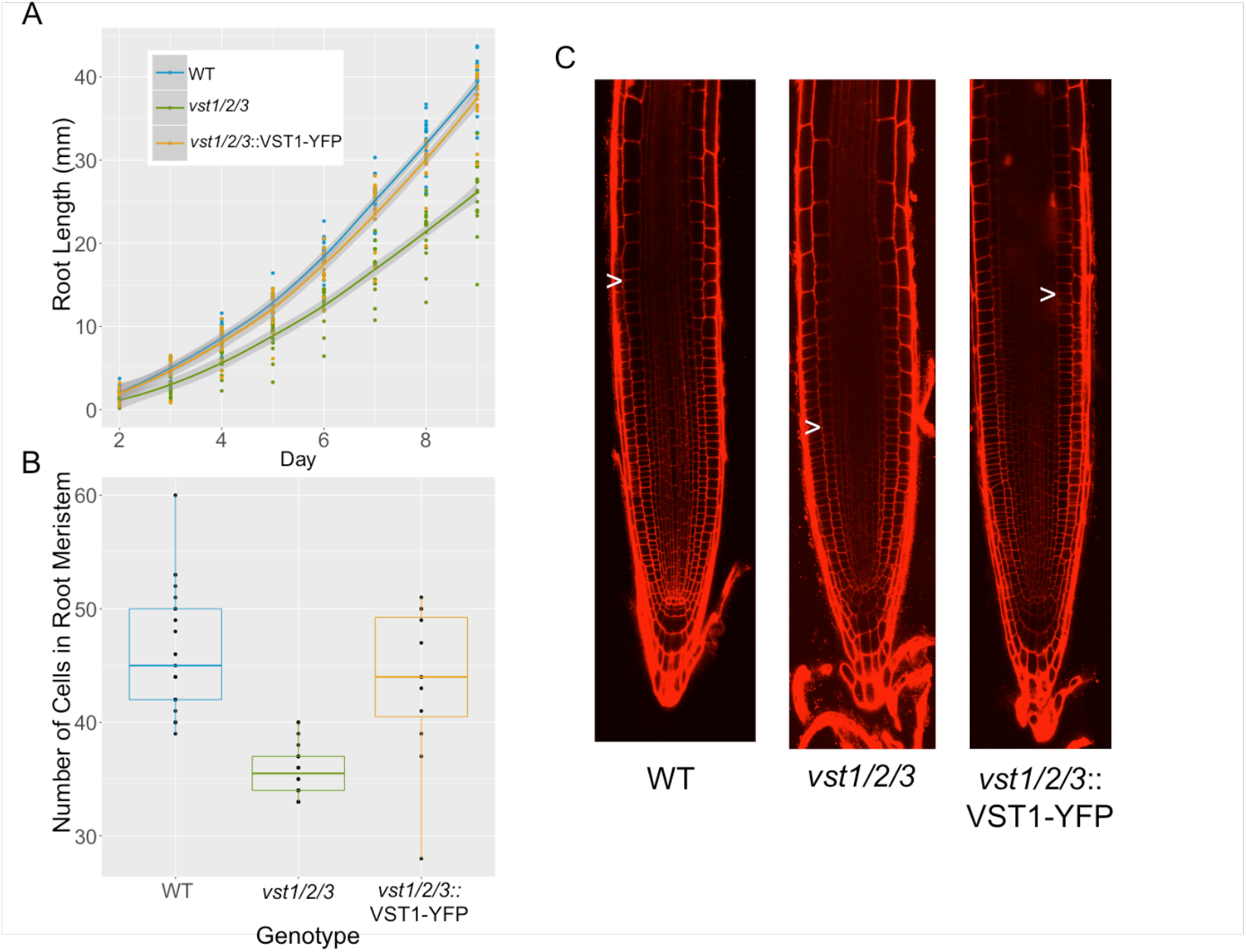
The reduced root length and meristem size in *Arabidopsis vst1/2/3* triple mutants is rescued by expression of *AtVST1*. (A) Timecourse of growth of *Arabidopsis* roots on agar plates. (B) Quantification of root meristem cell number and (C) representative confocal images of 7-day old *Arabidopsis* roots. Arrow indicates location of junction of meristem and elongation zone.

### *OsVST1* Can Complement the *Arabidopsis* VST Mutant Root Length and Aerial Phenotypes

We hypothesized that *OsVST1* might function in a similar manner as that of the *Arabidopsis* VSTs. To determine if *OsVST1* can complement the *Arabidopsis vst1/2/3* triple mutant we expressed *OsVST1* in the *vst1/2/3* background and found an increase in root length and plant height (Fig S9). These results indicate that *OsVST1* can functionally complement these *Arabidopsis* triple mutant phenotypes. This provides support to the hypothesis that OsVST1 may similarly act at PM-ER contact sites to facilitate signaling.

### *OsVST1* Mutant Tested in Field Conditions Shows Stable Grain Yield Properties

Using hydroponic and gel-based imaging systems, we found that *OsVST1* is required for wild-type RSA. Many RSA genes that have been cloned in monocot crops have highly pleiotropic phenotypes that would result in significant yield drag (Wissuwa et al., 2016). Our initial characterization of the *OsVST1* mutant indicated that its alleles have modest effects on shoot architecture (Fig S1D). This suggests that varying expression of *OsVST1* may be a means of modulating root architecture in the field without encountering a strong yield penalty.

We tested the E104-SR mutant in field trials in southern China. We found that plant height was shorter, yield per plant and tiller number were unaffected, but 1000-grain weight and seed setting rate are significantly higher in the mutant (Fig S10). This could reflect less energy put into root elongation that can be used on seed traits.

## Discussion

Using two different non-destructive imaging platforms we identified a novel regulator of RSA. We mapped the causal lesion underlying the E104-SR mutant root phenotype in an EMS-treated rice population to an 11-kilobase deletion (Fig S2). This is surprising considering the strong bias of EMS for transition mutations (Greene et al., 2003; Henry et al., 2014). However, larger deletions have been identified as likely causal mutations in EMS mutagenized populations in maize and wheat (Mo et al., 2018; Okagaki et al., 1991).

The *OsVST1* gene encodes a protein with high sequence similarity to the *Arabidopsis* VSTs. Coupled with the complementation of the root length and aerial phenotypes of *vst1/2/3* by *OsVST1*, this points to a potential role of this protein in regulating ER-PM contact sites. ER-PM interactions have been implicated in processes other than developmental signaling, including response to stress and establishment of cell to cell communication through plasmodesmata (Bayer et al., 2017; Lee et al., 2019)

Important open questions remain concerning the specific molecular pathways through which OsVST1 regulates RSA. OsVST1 could influence meristem size through broad modulation of cell-cell communication or through specific signaling complexes. In the context of *Arabidopsis* stomatal patterning, signaling through complexes specifically involving the ERECTA family of RLKs is affected in VST mutants (Ho et al., 2016). Considering the large number of RLKs that could control root meristem growth, it will be challenging to determine the specific signaling complexes affected in *OsVST1* mutants. The similarity of root meristem phenotypes between *Arabidopsis* and rice VST mutants may allow for use of this more genetically tractable system in understanding the molecular pathways affected by VST proteins in roots.

We observed a subcellular localization pattern of OsVST1 that is consistent with dual localization of the protein to both the nucleus and plasma membrane (Fig 4G). While any significance of this is currently unclear, movement of OsVST1 from the plasma membrane to the nucleus could serve as a way of controlling growth, perhaps in response to stress or nutrient conditions. It is also possible that nuclear localized OsVST1 may play a role in regulating gene expression that is functionally distinct from the ER-PM pathway.

VST homologs in Poplar were identified as differentially expressed during cell proliferation and radial growth in wood formation (Hertzberg et al., 2001; Schrader et al., 2004). In a patent application (US20170349910A1), overexpression of these genes resulted in trees with increased biomass. Our field results suggest that modulation of *OsVST1* does not have a detrimental effect on yield (Fig S10). Thus, there may be field conditions in which the shallow root architecture of the *OsVST1* mutant may be agronomically beneficial.

## Methods

### Plant Materials and Growth Conditions

For hydroponic growth experiments, germinated seeds were sown on floating nets and grown in full-strength Kimura nutrient solution (pH, 5.6) as described previously (Chen et al., 2013). The seedlings were photographed at 7 days after germination and transferred to nutrient solution to monitor root growth over the entire experimental period. Nutrient solution was changed every 7 days. The phenotypic characterization of plants was performed in a growth room under 14-hour day length at 60% to 70% humidity and 30/22°C (day/night). The growth room uses bulb type light with a photon density of ~300 μmol m^−2^ s^−1^.

For gel imaging experiments, plants were grown in solidified Yoshida’s Nutrient Solution (pH 5.8). After two days of pregrowth on petri plates in the dark at 30°C, seedlings were transplanted to sterile glass jars containing media. Plants were largely shielded from light and grown in a growth chamber under 12-hour day length at 30/27°C.

For field experiments, plants were grown at the Zhejiang University Experimental station in Hainan Province, China. Seeds were sown in early December, seedlings were transplanted after a month and harvesting and agronomic trait analysis was performed after maturation in mid-April of the next year.

### Mutagenesis and Screening

Seeds of rice cultivar Xiushui 134 (XS134) were treated with 0.8% ethyl methanesulfonate (EMS) for 10 hours at 25°C. Families of M2 plants were screened to identify lines with aberrant root length phenotypes. E104-SR was identified as a mutant segregating for a shorter root length compared to wild type.

### Gel-based Imaging and Analysis

Gel-grown plants were imaged daily using a previously described imaging system (Topp et al., 2013). 20-40 images were taken for each plant at each timepoint. Images were analyzed with the GiaRoots software (Galkovskyi et al., 2012).

### Mutation Mapping

The E104-SR mutant was backcrossed to the unmutagenized XS134 parent and the resulting F_1_ plant was selfed to produce an F_2_ mapping population segregating for the shorter root phenotype. Over 120 F_2_ plants were screened using the gel imaging system. Twenty plants with the shortest and most compact root systems were selected as the mutant class, while the twenty plants with the largest root systems made up the wild-type class.

DNA was isolated from leaf tissue from these individuals and the parental line XS134 using the Qiagen DNAeasy Plant Kit. These were collected in equal concentrations to make up the mutant bulk, wild-type bulk, and the XS134 sample. Library preparation and paired-end sequencing with the Illumina HiSeq 2500 were done at the Duke University Genomics Core Facility.

Reads were mapped to the Nipponbare (Oryza_sativa.IRGSP-1.0.31) reference genome using the “aln” function in BWA (Li & Durbin, 2009). SNPs and small indels were called using the “mpileup” function in SAMTools (Li, 2011). Allele counts we extracted from the resulting VCF file using an awk script from (Mascher et al., 2014). Background SNPs (also present in the XS134 sequence) were removed. The final VCF file was imported into the QTLseqr R package (Mansfeld & Grumet, 2018) and the “deltaSNP” index was calculated and plotted.

Mapped sequence reads were visualized with the Integrated Genome Browser (Freese et al., 2016).

### Vectors Construction and Transformation

CRISPR-Cas9 mediated gene editing was used to generate additional mutant alleles of *LOC_Os08g04620* and *LOC_Os08g04630.* Callus from XS134 was transformed with agrobacterium strain EHA105 containing constructs based on the vector pYLCRISPR/Cas9-MH containing gRNAs targeting exons of *LOC_Os08g04620* or *LOC_Os08g04630*. Diagrams are in Figures S3 and S4. Deletions were confirmed by Sanger sequencing. Two independent loss-of-function mutants were identified and phenotyped for each candidate gene.

Callus from E104-SR was transformed with agrobacterium strain EHA105 containing pCAMBIA1300 constructs with the *LOC_Os08g04620* (an 6896-bp DNA fragment containing a 3144-bp sequence upstream of the start codon, the entire gene sequence, and a 1651-bp sequence downstream of the stop codon) or *LOC_Os08g04630* (a 9466-bp DNA fragment containing a 3113-bp sequence upstream of the start codon, the entire gene sequence, and a 3199-bp sequence downstream of the stop codon) open reading frames. Two independent potential complementation lines were identified and phenotyped for each candidate gene. To construct the promoter: GUS reporter lines, callus from XS134 was transformed with agrobacterium strain EHA105 containing pCAMBIA1300 construct with *GUS* reporter gene driven by the *LOC_Os08g04620* promoter (the 3139-bp sequence upstream of the start codon).

To construct the OsVST1 and GFP fusion protein lines. Two constructs of *ProVST1:VST1:GFP* (*VST1:GFP*) and *ProVST1:GFP:VST1* (*GFP:VST1*) were generated. A 5237-bp DNA fragment containing genomic *OsVST1* and the 3139-bp region upstream of the *OsVST1* start codon fused in-frame to the 5’ end of GFP in the modified pCAMBIA1300-GFP vector to generate *ProVST1:VST1:GFP*. A 3747-bp DNA fragment containing genomic *OsVST1* and the 1646-bp sequence downstream of the stop codon fused in-frame to the 3’ end of GFP, and driven by the *OsVST1* promoter (the 3139-bp sequence upstream of the start codon) to generate *ProVST1:GFP:VST1*. Callus from E104-SR was transformed with agrobacterium strain EHA105 containing *ProVST1:VST1:GFP* or *ProVST1:GFP:VST1* constructs. The primers used (including the restriction enzyme sites used in the cloning) are listed in Supplemental Table 1.

### Protein Sequence Alignment

Proteins containing a Major Sperm Protein Domain were downloaded from the MSU Rice Genome Database and The Arabidopsis Information Resource. Proteins encoding the longest isoforms of each gene were used. Sequences were aligned using MAFFT (Madeira et al., 2019).

### Root Staining and Fluorescence Imaging

Seedlings were stained with beta-galactosidase solution for 6 h at 37°C, and then de-colored with ethanol for two days to remove chlorophyll. Images of seedling or root were taken using a camera (Nikon) or stereomicroscope (Leica), respectively. To observe meristem and elongation zone of root tip, rice root tips were fixed in 2.5% glutaric dialdehyde fixation solution overnight, dehydrated using acetone, infiltrated and embedded in Spurr’s resin, and then sectioned into 2 μm sections. Sections were mounted on slides and stained with methylene blue solution. Images were taken using a microscope (Nikon).

For the GFP fluorescence analysis, rice roots were embedded using 4% agarose. Images of cross- or longitudinal-sections of root were taken using a LSM710 confocal laser scanning microscope (Zeiss) after cutting the root into 40- to 80- μm sections using a microtome (Leica).

### Arabidopsis Root Growth Assays

For Arabidopsis experiments, seeds were surface sterilized with 50% bleach for 10 minutes, rinsed 5x with sterile water, then sown on ½ MS 1% sucrose plates solidified with 1% agar in square petri dishes, sealed with micropore tape, stratified at 4°C for 2 days in the dark before moving to a 22°C, 16 hour light cycle growth chamber vertically. Under these conditions germination occurred approximately 1 day after moving to the chamber. For analysis of root growth dynamics of WT Arabidopsis, the *vst1/2/3* triple mutant, and the *vst1/2/3 AtVST1-YFP* complemented line, plates were imaged every day between 2 and 9 post-germination, and root lengths for plants were measured in imageJ. In a separate experiment, cortical meristem size was measured for WT, *vst1/2/3*, and *vst1/2/3 AtVST1-YFP* 7 days after germination by staining with 10 mg /ml propidium iodide for approximately 2 minutes before imaging with a Zeiss 510 confocal microscope. Meristem cells were counted from the first cortical cell after the cortex/endodermal initial or quiescent center (in the case where the initial had asymmetrically divided) until the abrupt increase in cell length observed in the transition zone. Counting was performed in imageJ, and two independent double-blinded counts were performed with nearly identical results in order to validate the apparent difference in meristem size.

For the *OsVST1* complementation experiment, we synthesized the *OsVST1* cDNA sequence (GenScript) to include attL1/L2 sites for direct gateway Recombination recombination into pGWB502-omega (Nakagawa et al., 2007), which contains a 35S promoter and translational enhancer for overexpression. The *vst1/2/3* mutant was transformed by floral dip and transformants were selected on hygromycin. We selected two independent lines in the T2 that appeared to be segregating for an increase in plant height. We isolated homozygous lines for each in the T4 based on uniform hygromycin resistance. To test complementation of root length, we grew WT, *vst1/2/3,* and *vst1/2/3 OsVST1* in the same manner as above, and measured root length was measured at 8 days post-germination. Seedlings of these plants were transferred to soil and plant height measured approximately 5 weeks later.

## Accession Numbers

Sequence data from this article can be found in the NCBI SRA under Bioproject PRJNA631002

## Acknowledgements

The authors would like to thank Dr. Dominique Bergmann for the vst1/2/3 and complementation lines, Dr. Colleen Drapek for critical reading of this manuscript, members of the Benfey and Mao labs for helpful discussions, and Alan Tang, Medhavinee Mijar, and Trevor Gannalo for their help with plant care and image processing. Listing order for co-lead authors was determined by the last digit of the closing price (odd or even) of the Shanghai Stock Exchange Composite Index on May 12, 2020.

