## Supplemental Figures 1-10 for "VST Family Proteins are Regulators of Root System Architecture in Rice and *Arabidopsis*"

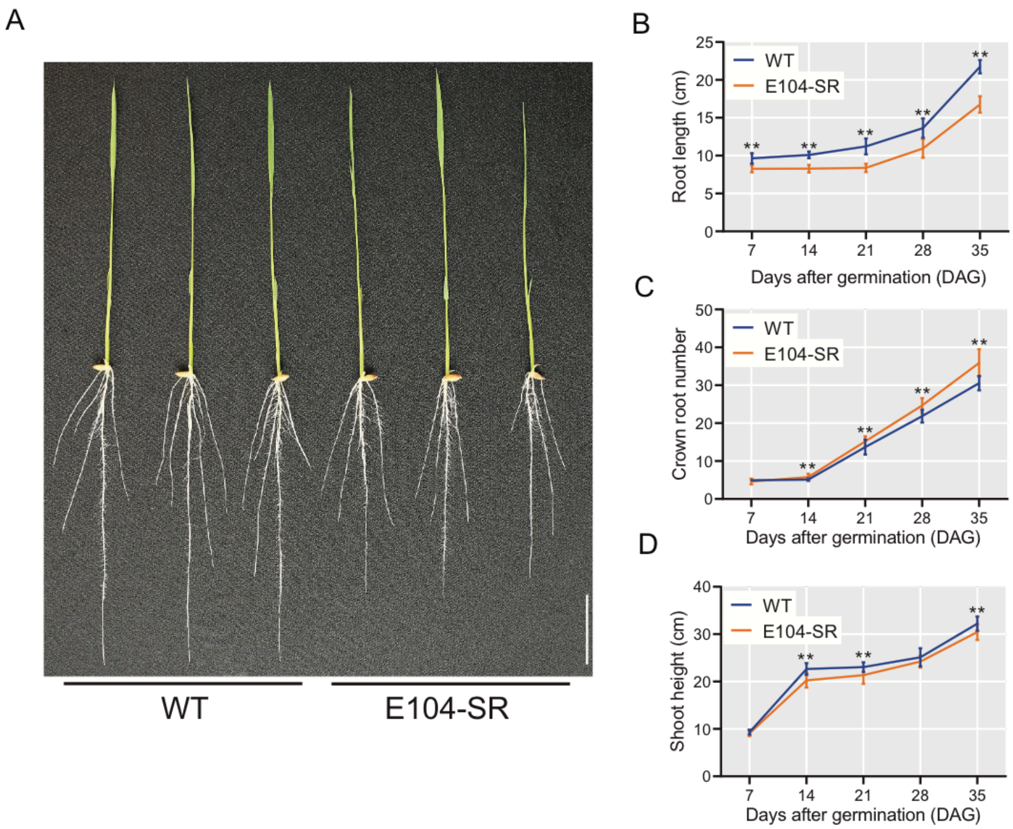


Figure S1: E104-SR is a mutant with decreased root length and slightly decreased shoot height. (A) Image of 7-day-old wild type and E104-SR plants, grown in hydroponic culture. Bar = 3 cm. (B) Root length of wild-type and E104-SR. (C) Crown root number of wild-type and E104-SR. (D) Shoot height of wild-type and E104-SR. Data represent means ± SD (*n* = 20). Data significantly different in E104-SR from wild type are indicated (**P < 0.01; Student's t test).

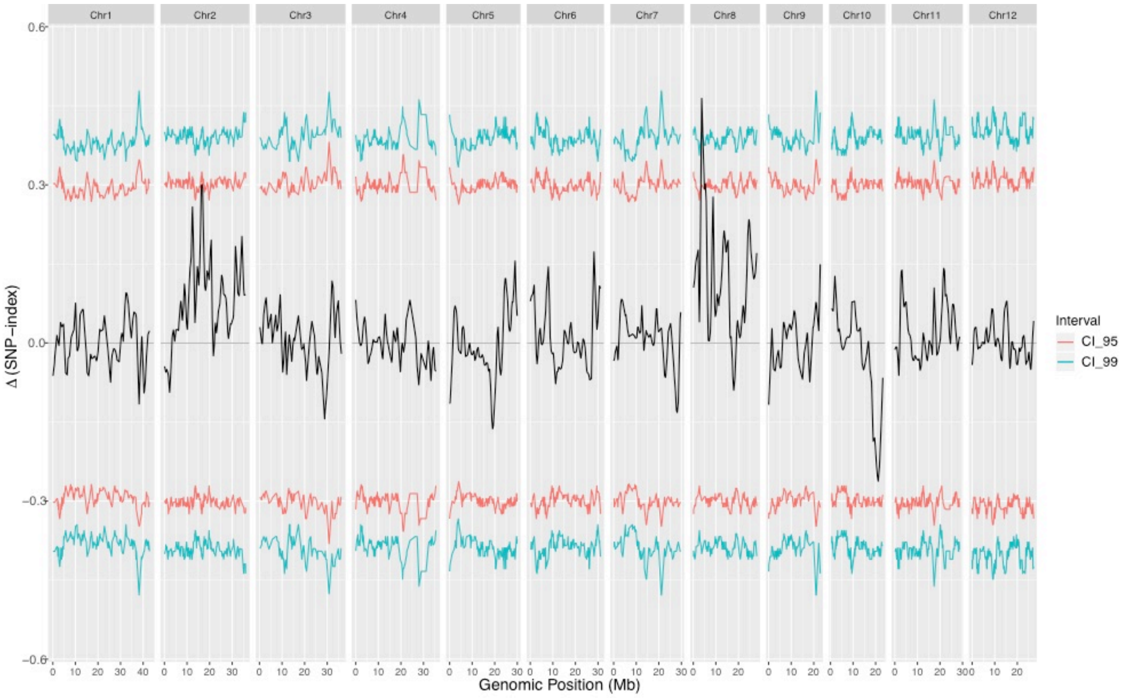


Figure S2: Bulked-segregant analysis identifies a peak on Chromosome 8 associated with the E104-SR root length phenotype. Allele frequencies of SNPs called from mutant and wild-type bulks were calculated, compared, and plotted using the R package QTLseqr v0.7.3 (Mansfeld & Grumet, 2018).

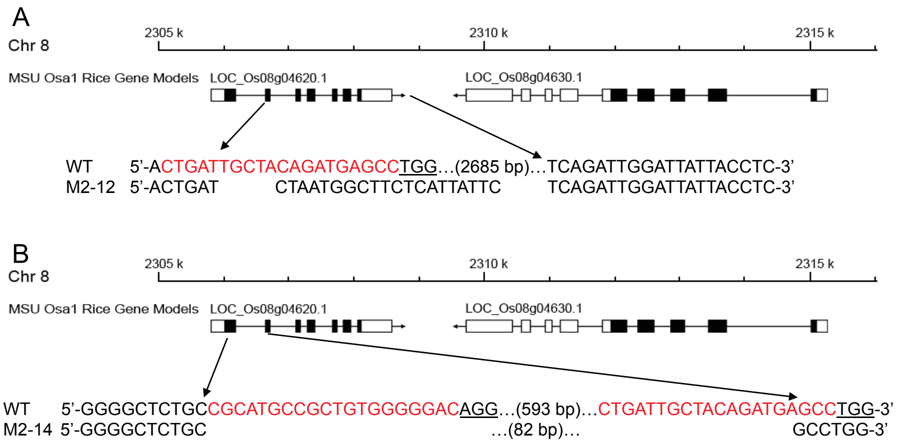


Figure S3: CRISPR-Cas9-mediated target mutagenesis of *LOC_Os08g04620*. Top: a schematic diagram of the *LOC_Os08g04620* gene. The lower panel shows alignment of XS134, M2-12 (A) and M2-14 (B) sequences containing the CRISPR-Cas9 target sites. The 20 bp CRISPR-Cas9 target sequences adjacent to the underlined protospacer adjacent motifs (PAMs) are indicated in red in XS134 sequences. The newly created M2-12 or M2-14 mutants contain a large substitution of 2703 bp to 20 bp or 633 bp to 82 bp, respectively. The interval shown by the arrow indicates the position of large substitution in *LOC_Os08g04620*.

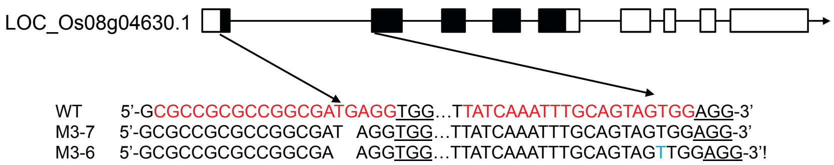


Figure S4: CRISPR-Cas9-mediated target mutagenesis of *LOC_Os08g04630*. Top: a schematic diagram of the *LOC_Os08g04630* gene. The lower panel shows alignment of XS134, M3-7 and M3-6 sequences containing the CRISPR-Cas9 target sites. The arrows indicate the position of CRISPR-Cas9 target sites in *LOC_Os08g04630* gene. The 19 or 20 bp CRISPR-Cas9 target sequences adjacent to the underlined protospacer adjacent motifs (PAMs) are indicated in red in XS134 sequences. The newly created M3-7 or M3-6 mutants contain deletions or insertions represented with black or blue letters, respectively.

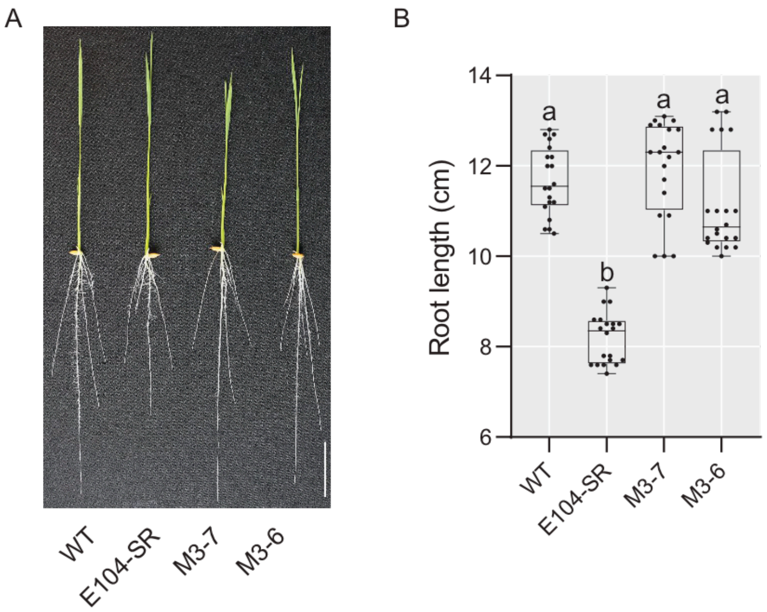


Figure S5: CRISPR-mediated deletion of *LOC_Os08g04630* does not result in decreased root length. (A) Phenotypes of 7-day-old knockout transgenic lines grown in hydroponic culture. Median is represented by horizontal line in the box plot, box ranges represent quartiles 1 and 3, and minimum and maximum values are represented by error bars ($n$ = 20). Significantly different values are indicated by different letters ($P < 0.01$; one-way ANOVA with Tukey's test).

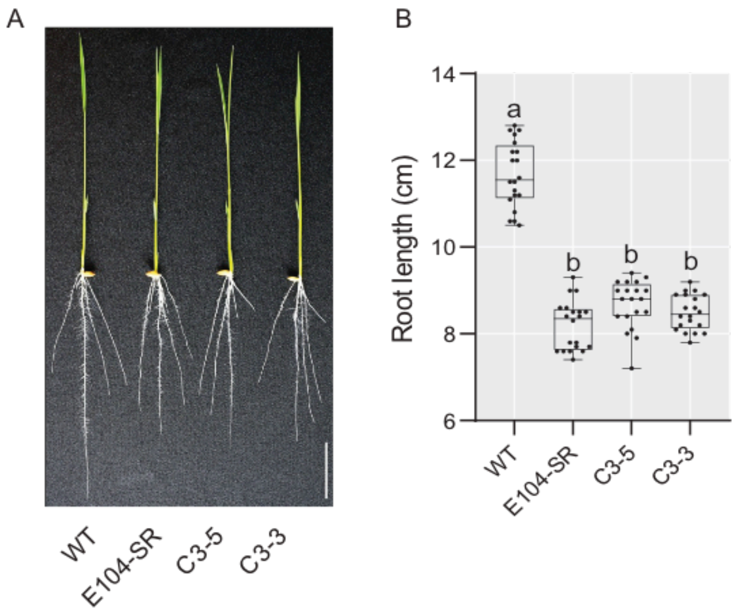


Figure S6: Insertion of genomic fragment containing *LOC_Os08g04630* into E104-SR does not complement the root length phenotype. (A) Phenotypes of 7-day-old complementation lines grown in hydroponic culture. C3-3 and C3-5 are two independent T2 transgenic lines of E104-SR containing a genomic fragment with only *LOC_Os08g04630*. Bar = 3 cm. (B) Root lengths of 7-day-old complementation lines grown in hydroponic culture. Median is represented by horizontal line in the box plot, box ranges represent quartiles 1 and 3, and minimum and maximum values are represented by error bars ($n = 20$). Significantly different values are indicated by different letters ($P < 0.01$; one-way ANOVA with Tukey's test).

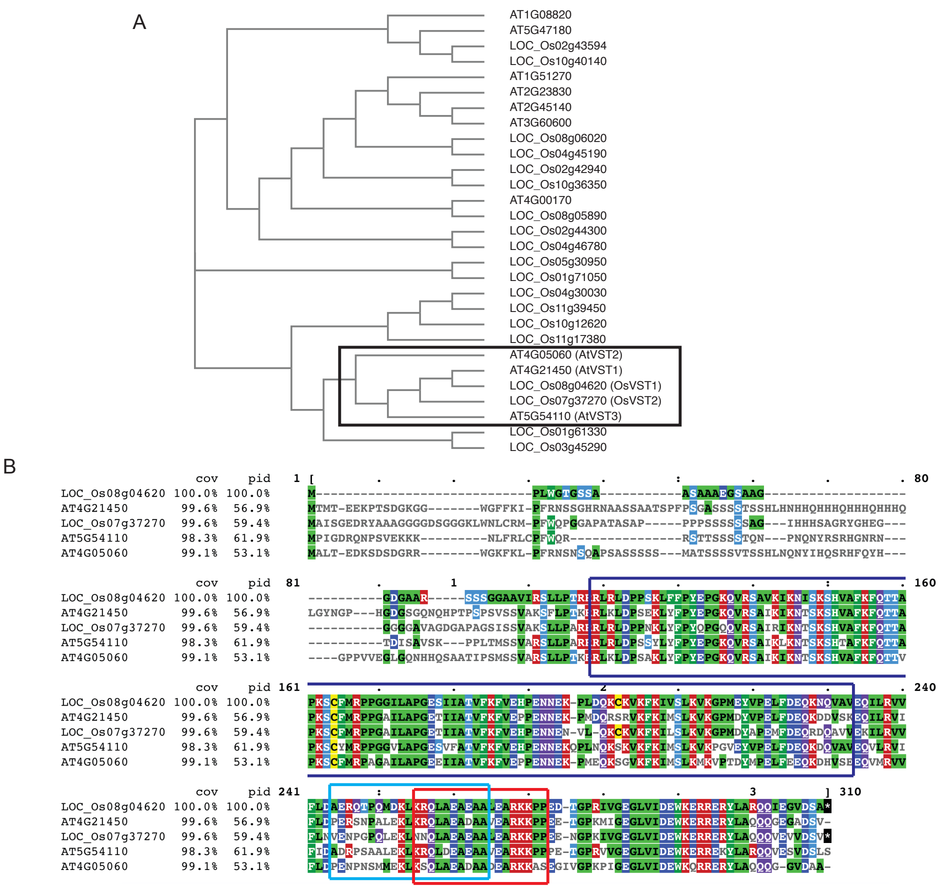


Figure S7: Sequence alignment of MSP domain-containing proteins in Rice and *Arabidopsis* shows clustering of LOC_Os08g04620 with AtVST proteins. (A) Cladogram of MAFFT sequence alignment of *Arabidopsis* and rice MSP-domain containing proteins. (B) MAFFT sequence alignment of *Arabidopsis* and rice VST proteins. Purple, light blue, and red boxes indicate locations in LOC_Os08g04620 of predicted MSP domain, coiled-coil domain, and nuclear localization sequences, respectively.

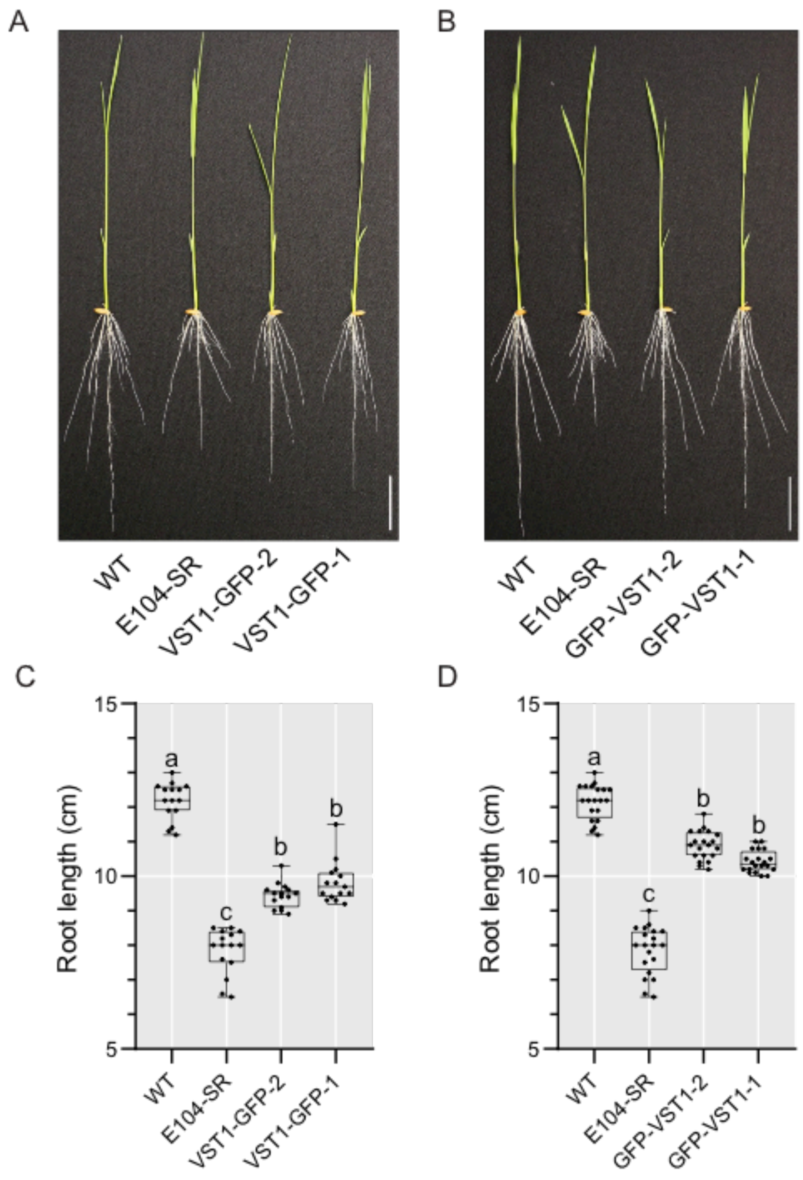


Figure S8: *GFP-VST1* complements the short root phenotype of E104-SR better than *VST1-GFP*. (A) Phenotype of two *VST1-GFP* (*VST1-GFP-1* and *VST1-GFP-2*) transgenic lines at seven days. Bar = 3 cm. (B) Phenotype of two *GFP-VST1* (*GFP-VST1-1* and *GFP-VST1-2*) transgenic lines at seven days. Bar = 3 cm. (C) Root length of wild type, E104-SR, and two *VST-GFP* transgenic lines. Median is represented by horizontal line in the box plot, box ranges represent quartiles 1 and 3, and minimum and maximum values are represented by error bars ($n$ = 15). Data significantly different from the corresponding plants are indicated by different letters ($P < 0.01$; one-way ANOVA with Tukey's test). (D) Root length of wild type, E104-SR, and two *GFP-VST1* transgenic lines. Median is represented by horizontal line in the box plot, box ranges represent quartiles 1 and 3, and minimum and maximum values are represented by error bars ($n$ = 20). Data significantly different from the corresponding plants are indicated by different letters ($P < 0.01$; one-way ANOVA with Tukey's test).

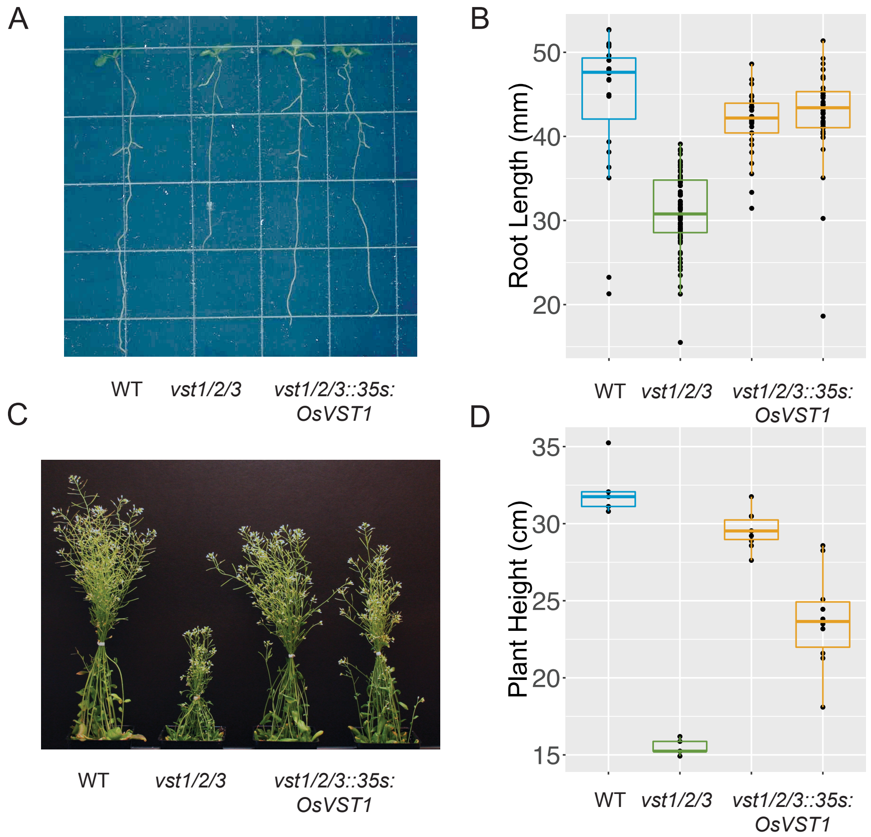


Figure S9: Expression of *OsVST1* can partially rescue the decreased root length and plant height phenotypes in the *Arabidopsis vst1/2/3* triple mutant. Representative root growth (A) and quantification of root growth (B) on agar plates. Representative plants (C) and quantification of plant height (D) at approximately six weeks post planting. Replicates of complementation lines are from two independent transformants.

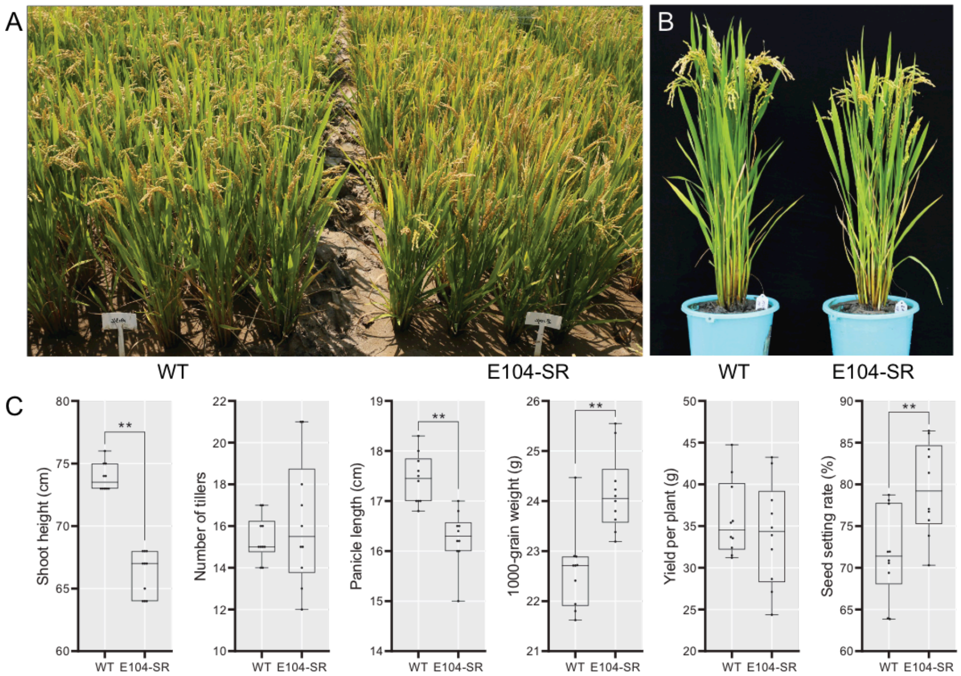


Figure S10: E104-SR has decreased plant height and slightly increased 1000-grain weight when grown in field conditions. (A) and (B) Morphological phenotypes of wild type and E104-SR growing in Hainan province. (C) Statistical analysis of agronomic traits of wild type and E104-SR growing field. Median is represented by horizontal line in the box plot, box ranges represent quartiles 1 and 3, and minimum and maximum values are represented by error bars ($n$ = 10). ** Significant difference between XS134 and E104-SR ($P < 0.01$; Student's t test); * Significant difference between XS134 and E104-SR ($P < 0.05$; Student's t test).
