## Supplemental Table 1 for "VST Family Proteins are Regulators of Root System Architecture in Rice and *Arabidopsis*"

Supplemental Table 1. Sequence of the Primers Used in This Study.

|  | Primer Name | Sequences (5’-3’) |
| --- | --- | --- |
| For E104-SR Mutant and Wild Type Identification | E104-SR-F | GTTAATTGCCCTCGATCCAAAC |
|  | E104-SR-R | TCGGTATCAATCACATGTCTCC |
|  | XS134-F | GAAGGTAGGAGGAGTAGAGAGAAG |
| For qRT-PCR | qPCR-VST1-F | CAAGATCAAGAACATCAGCAAGTC |
|  | qPCR-VST1-R | TCTCAGGATGCTCCACAAAC |
| Primers for Vector Construction |  |  |
| LOC_Os08g04620-CRISPR Target | LOC_Os08g04620-U3-F | GGCACGCATGCCGCTGTGGGGGAC |
|  | LOC_Os08g04620-U3-R | AAACGTCCCCCACAGCGGCATGCG |
|  | LOC_Os08g04620-U6a-F | GCCGCTGATTGCTACAGATGAGCC |
|  | LOC_Os08g04620-U6a-R | AAACGGCTCATCTGTAGCAATCAG |
| LOC_Os08g04630-CRISPR Target | LOC_Os08g04630-U3-F | GGCATATCAAATTTGCAGTAGTGG |
|  | LOC_Os08g04630-U3-R | AAACCCACTACTGCAAATTTGATA |
|  | LOC_Os08g04630-U6a-F | GCCGCGCCGCGCCGGCGATGAGG |
|  | LOC_Os08g04630-U6a-R | AAACCCTCATCGCCGGCGCGGCG |
| LOC_Os08g04620 Complementation | LOC_Os08g04620-SalI-F | GCGTCGACAGCTATCACGAGCAGACTCCATTTC |
|  | LOC_Os08g04620-BamHI-R | CGGGATCCGACCAATAGTTTGTCTTTGCAGTTC |
| LOC_Os08g04630 Complementation | LOC_Os08g04630-infusion-F1 | AATTCGAGCTCGGTACCCGGGCCTCTACTTTCTCTCTCCTCTCTT |
|  | LOC_Os08g04630-infusion-F1 | GTAGGAGGAGTAGAGAGAAGGTGGTGGAGGAGGAGGAGAC |
|  | LOC_Os08g04630-infusion-F2 | CTTCTCTCTACTCCTCCTACCTTC |
|  | LOC_Os08g04630-infusion-F2 | GACCAAACAGCCATTCCATAAG |
|  | LOC_Os08g04630-infusion-F3 | TATGGAATGGCTGTTTGGTCCACTGGTATTGGTACCCGGCCCTT |
|  | LOC_Os08g04630-infusion-F3 | CTTGCATGCCTGCAGGTCGACCCTGTGTGACCTGTACTGATATAAT |
| *ProVST1:GUS* | ProVST1-SalI-F | CTTGCATGCCTGCAGGTCGACTCACGAGCAGACTCCATTTC |
|  | ProVST1-XbaI-R | TGGTACCGTGGATCCTCTAGAGCGGCAGAGCCCCAAGGTAC |
| *ProVST1:VST1:GFP* | ProVST1-SalI-F | CTTGCATGCCTGCAGGTCGACTCACGAGCAGACTCCATTTC |
|  | gVST1-BamHI-R | TCACCATGGTACCGTGGATCCTCGCTGAATCAACCCCTTCAATCT |
| *ProVST1:GFP:VST1* | ProVST1-infusion-F1 | GAGCTCGGTACCCGGGGATCCTCACGAGCAGACTCCATTTC |
|  | ProVST1-infusion-R1 | AAGGTACCCTTGTACAGCTCGTCCATGCCGTGA |
|  | gVST1-infusion-F2 | GTACAAGGGTACCTTGGGGCTCTGCCGCAT |
|  | gVST1-infusion-R2 | CTTGCATGCCTGCAGGTCGACATAGTTTGTCTTTGCAGTTC |
